## Supplemental files for "Massively parallel jumping assay decodes *Alu* retrotransposition activity"

### Extended Data Figures and legends

a

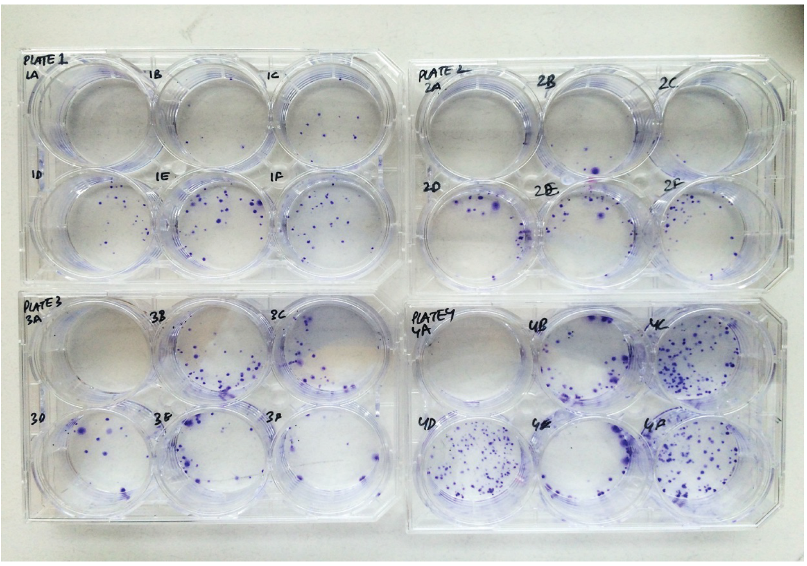


b

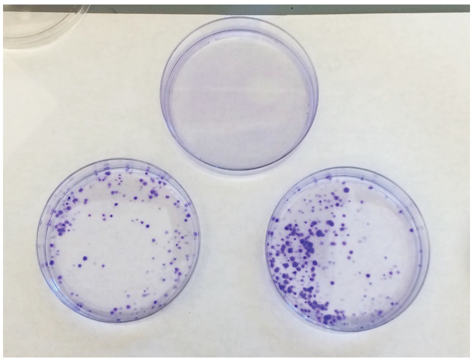

**Extended Data Fig 1**: **Retrotransposition colony assay.**
Plasmids were transfected in Hela cells and selected on G-418 for 3-4 weeks to obtain colonies that were stained with Crystal Violet staining solution.
**a**, Optimization of positive control element AluYa5 with two different Transposase helper plasmids L1-helper and ORF2 helper transfections at different concentrations. Plate 1 (top left) Well A (top left in plate) (negative control) contains 500ng of AluYa5 transfected without a L1 helper. Wells B-F have 500ng of AluYa5 transfected along with increasing concentrations L1-helper plasmid: 100ng (1B), 200ng (1C), 300ng (1D), 400ng (1E), 500ng (1F). Plate 2 (top right) Well A (top left in plate) (negative control) has 1000ng of AluYa5 transfected without L1 helper. Wells B-F have 1000ng of AluYa5 transfected along with increasing concentrations L1-helper plasmid: 100ng (2B), 200ng (2C), 300ng (2D), 400ng (2E), 500ng (2F) respectively. Plate 3 (bottom left) Well A (top left in each plate) (negative control) has 500ng of AluYa5 transfected without an ORF2 helper. Wells B-F have 500ng of AluYa5 transfected along with increasing concentrations ORF2 helper plasmid: 100ng (3B), 200ng

(3C), 300ng (3D), 400ng (3E), 500ng (3F) respectively. Plate 4 (bottom right) WellA (top left in each plate) (negative control) has 1000ng of AluYa5 transfected without an ORF2 helper. Wells B-F have 1000ng of AluYa5 transfected along with increasing concentrations ORF2 helper plasmid: 100ng (4B), 200ng (4C), 300ng (4D), 400ng (4E), 500ng (4F) respectively.
**B**, Colony assay optimization at larger scale in 15mm dishes. The top plate is a negative control containing transfections with 2.5ug of AluYa5 alone (no ORF2 control), the bottom left plate was transfected with 2.5ug of AluYa5 and 10ug of ORF2 helper plasmids, and the bottom right plate was transfected with 5ug of AluYa5 and 10ug of ORF2 helper plasmids.

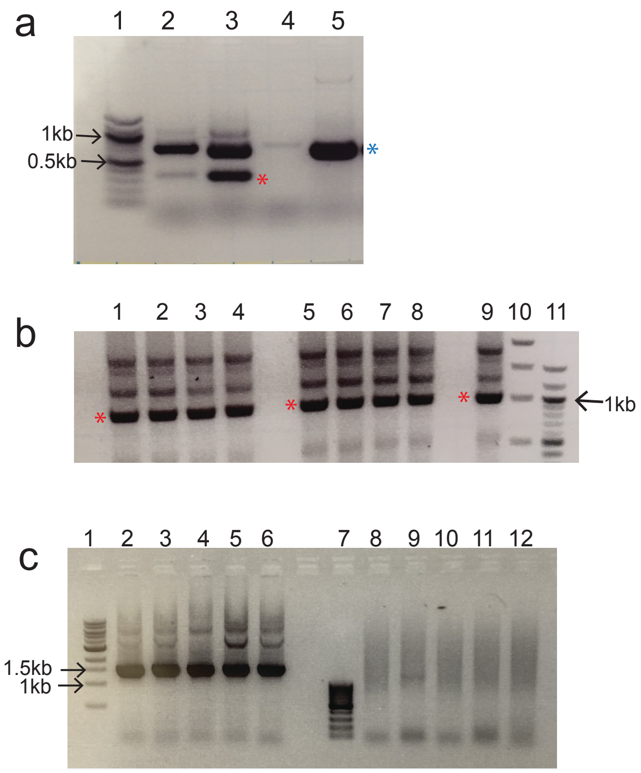


**Extended Data Fig. 2: Alu-Neo cassette amplification to differentiate between retrotransposition vs random integration events.**
**a**, Agarose gel showing that the Alu-Neo cassette in the plasmid with the intron is around 800bp (Lane5, blue asterisk) while the spliced cassette is around 330bp (Lane 3 red asterisk) using primers NeoEx1F ATGGGATCGGCCATTGAACAAGATG and NeoEx2R GCAAGGTGAGATGACAGGAGATCC. Lane 1, DNA ladder. Lane 2, PCR on genomic DNA isolated from AluYa5+L1 helper transfections, Lane 3 PCR on genomic DNA isolated from AluYa5+ORF2 helper transfections, Lane 4 no DNA, Lane5, PCR amplification of Alu-Neo cassette from AluYa5 plasmid.
**b**, Agarose gel showing the retreival of Alu-Neo cassette (spliced, 1kb red asterisk) from HeLa cells transfected with Alu6b, (Lanes,1 and 5), Alu14b (Lanes 2 and 6), Aluh1.1 (Lanes 3 and 7), AluSx (Lanes 4 and 8), AluYa5 (Lane 9) using primer sets described in **Supplementary Table 5**). DNA ladders (1kb and 0.1kb ladders NEB Cat no: N0468S and N3231S respectively) are in lane 10 and 11.
**c**, Agarose gel confirming that there is no amplification from transfected HeLa cells cells (Lanes 8-12) using the same primer sets above in (**b**) compared to 1.5kb amplicon from plasmids AluSx, Alu14b, AluYa5, Alu6b, Aluh1.1 from Lanes 2-6 respectively. The DNA ladder (1kb ladders NEB Cat no: N0468S) is in Lane1.

a

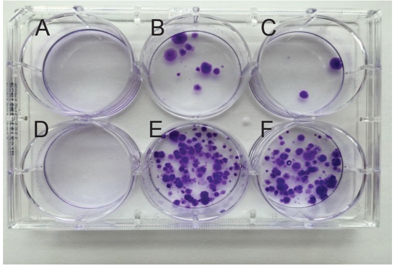


b

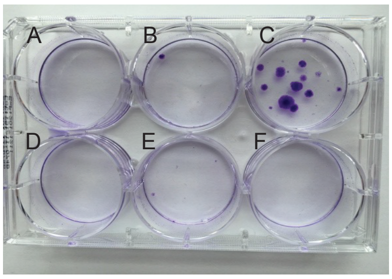


**Extended Data Fig. 3: Small scale retrotransposition colony assay with Alu mutagenized library.**

Retrotransposition plasmids were transfected in Hela cells and selected on G-418 for 3-4 weeks to obtain colonies that were stained with Crystal Violet staining solution.

**a**, Small scale colony assay optimization with a mutagenized Alu library with ORF2 Transposase helper plasmid. Well A (top left in plate) is a negative control that has 1ug of Alu6b transfected without a ORF2 helper. Wells B-C have 1ug of Alu6b transfected and 1ug of Mut-Alu6b along with 1ug of ORF2-helper plasmid. Well D (bottom left) in a negative control that has AluSx without any ORF2. Wells E-F have 1ug of AluSx and 1ug of Mut-AluSx respectively along with 1ug of ORF2 helper plasmids.

**b**, Small scale colony assay optimization with mutagenized Alu library with ORF2 Transposase helper plasmid. Well A (top left in plate) is a negative control that has 1ug of Aluh1.1 transfected without any ORF2 helper.  Wells B-C have 1ug of Aluh1.1 transfected and 1ug of Mut-Aluh1.1 along with 1ug of ORF2-helper plasmid. Well D (bottom left) in a negative control that has Alu14b without an ORF2. Wells E-F have 1ug of Alu14b and 1ug of Mut-Alu14b respectively along with 1ug of ORF2 helper plasmids.

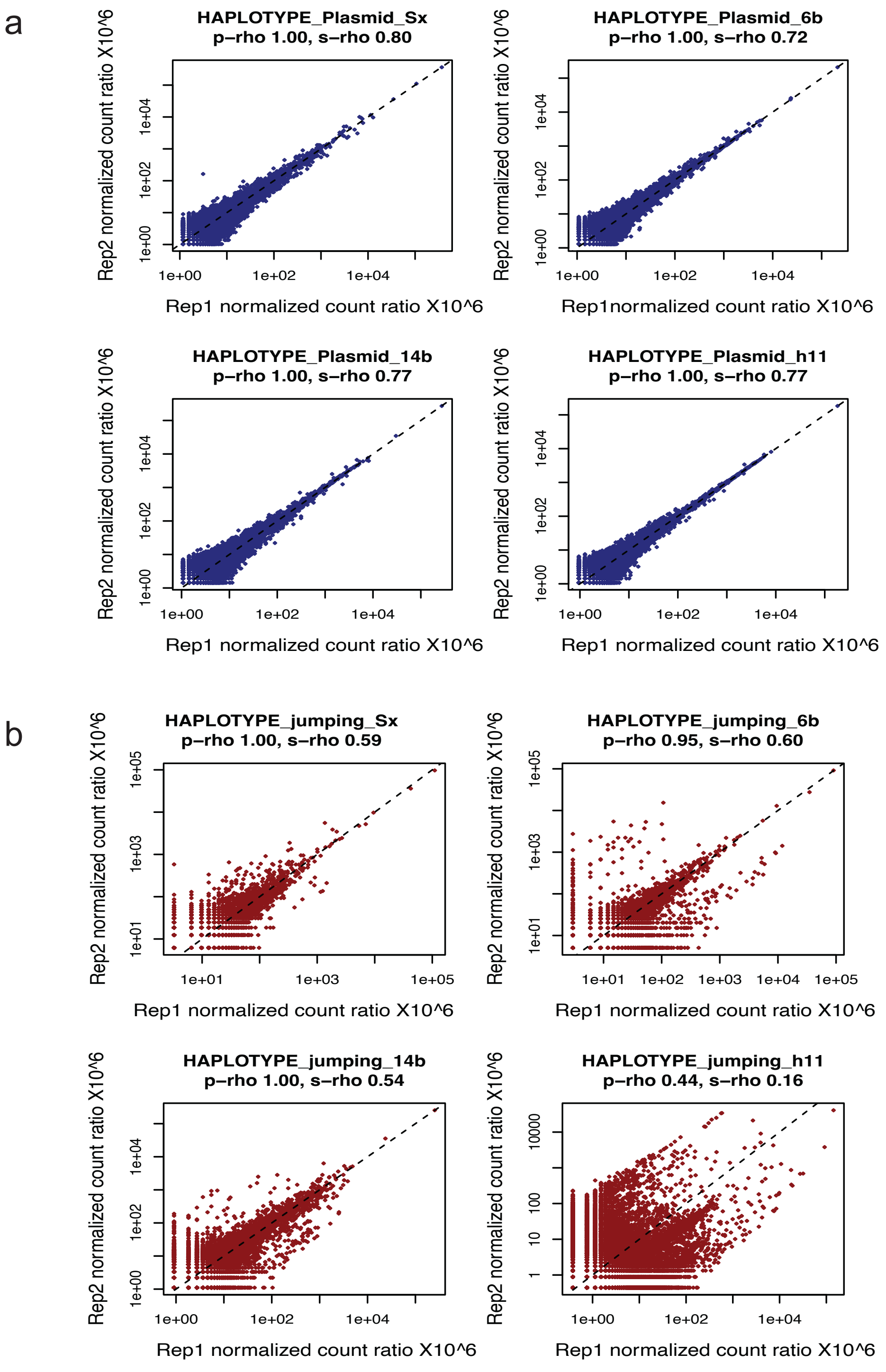


**Extended Data Fig. 4: Scatter plots showing correlations for the unique variants or haplotypes detected in each replicate.**

**a-b**, Top panel with blue dots (**a**) depicts the correlation between plasmid mutagenesis libraries and the bottom panel with red dots (**b**) shows the correlation between two independent jumping experiments with mutagenized libraries.

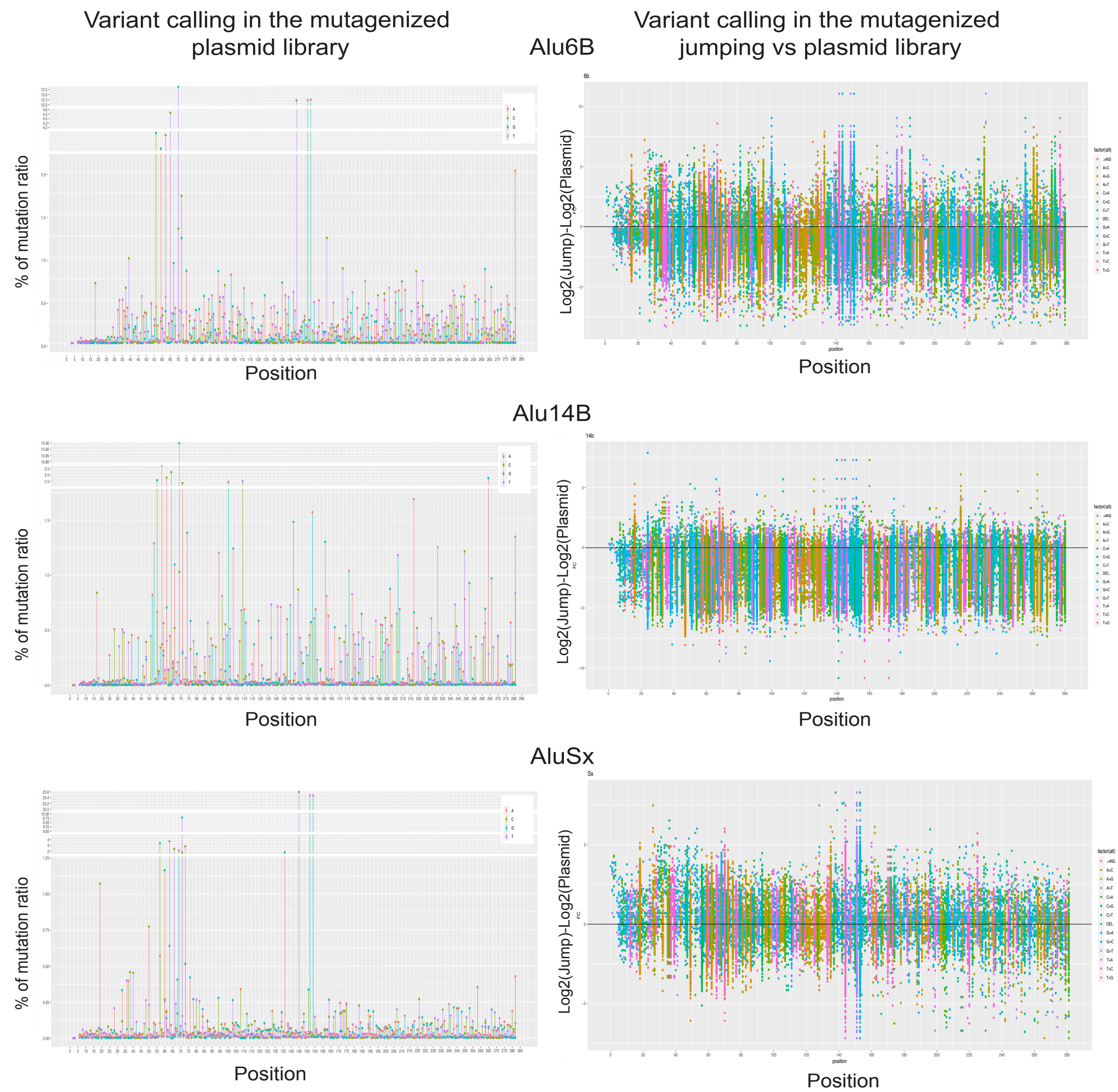


**Extended Data Fig. 5: Variants in the libraries for each nucleotide position in Alu.**
Left panel: Lollipop plots for variants at each position along the length of Alu element found in the plasmid mutagenesis library. The variants with high percentage of occurrence in the library could be caused due to founder effect of certain variants getting fixed in early PCR cycle. Right panel: Density plot of nucleotide changes (transitions, transversions, insertion or deletion) at each position detected in both plasmid and jumping library with Log2FC of jumping vs plasmid normalized at 0Log2FC to be the reference sequence.

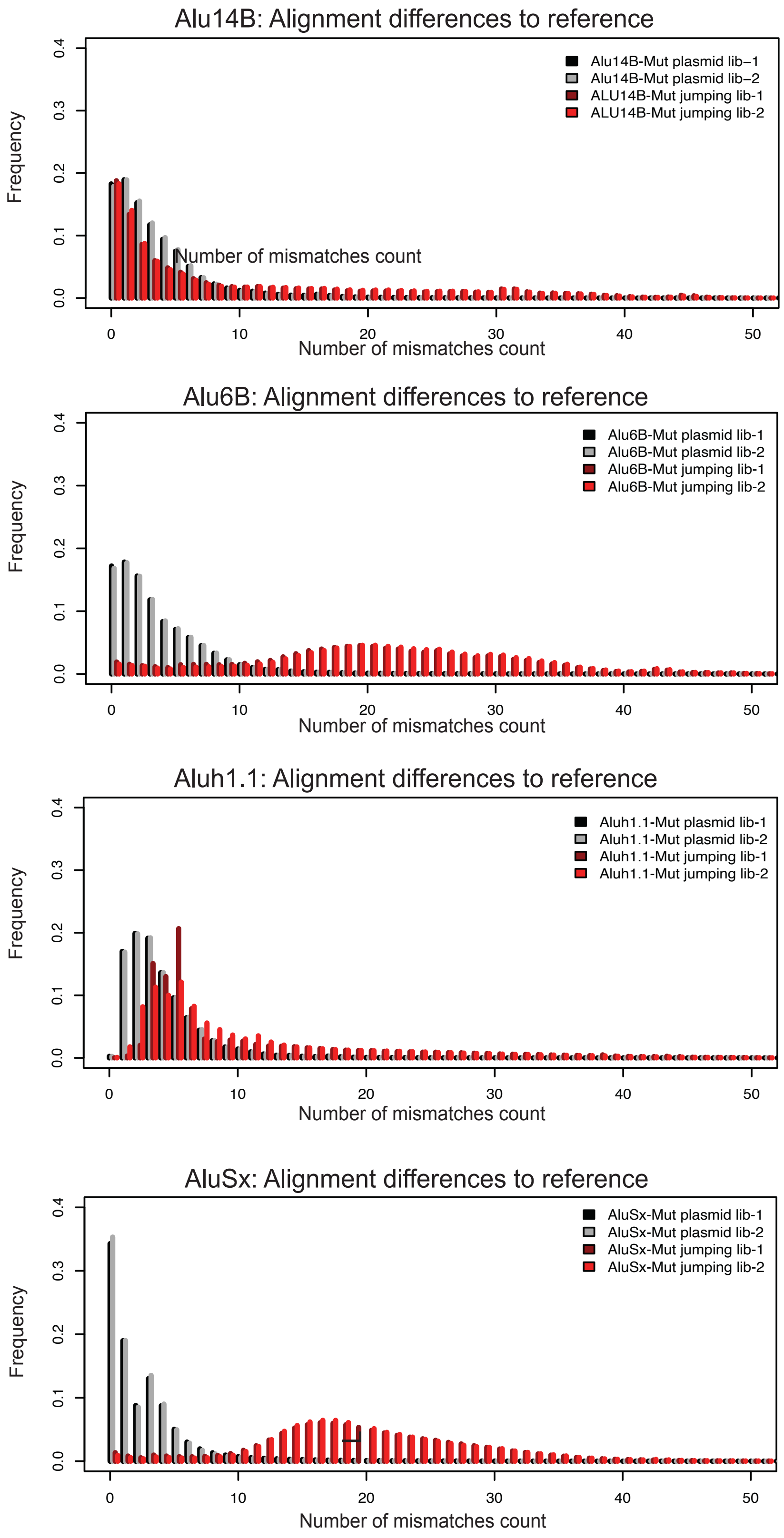


**Extended Data Fig. 6: Alignment of mutagenized Alu library to its reference sequence.** Frequency of reads with stratified number of mismatches on the X axis. Number of the mismatches is calculated by the NM tags, NM-Edit distance tag records the Levenshtein distance between the read the reference. Up to a 10-mismatch cutoff was found to be sufficient to incorporate in the analysis. In some of the jumping libraries, there are some reads with an unexpectedly high number of mismatches that were not considered for further analysis.

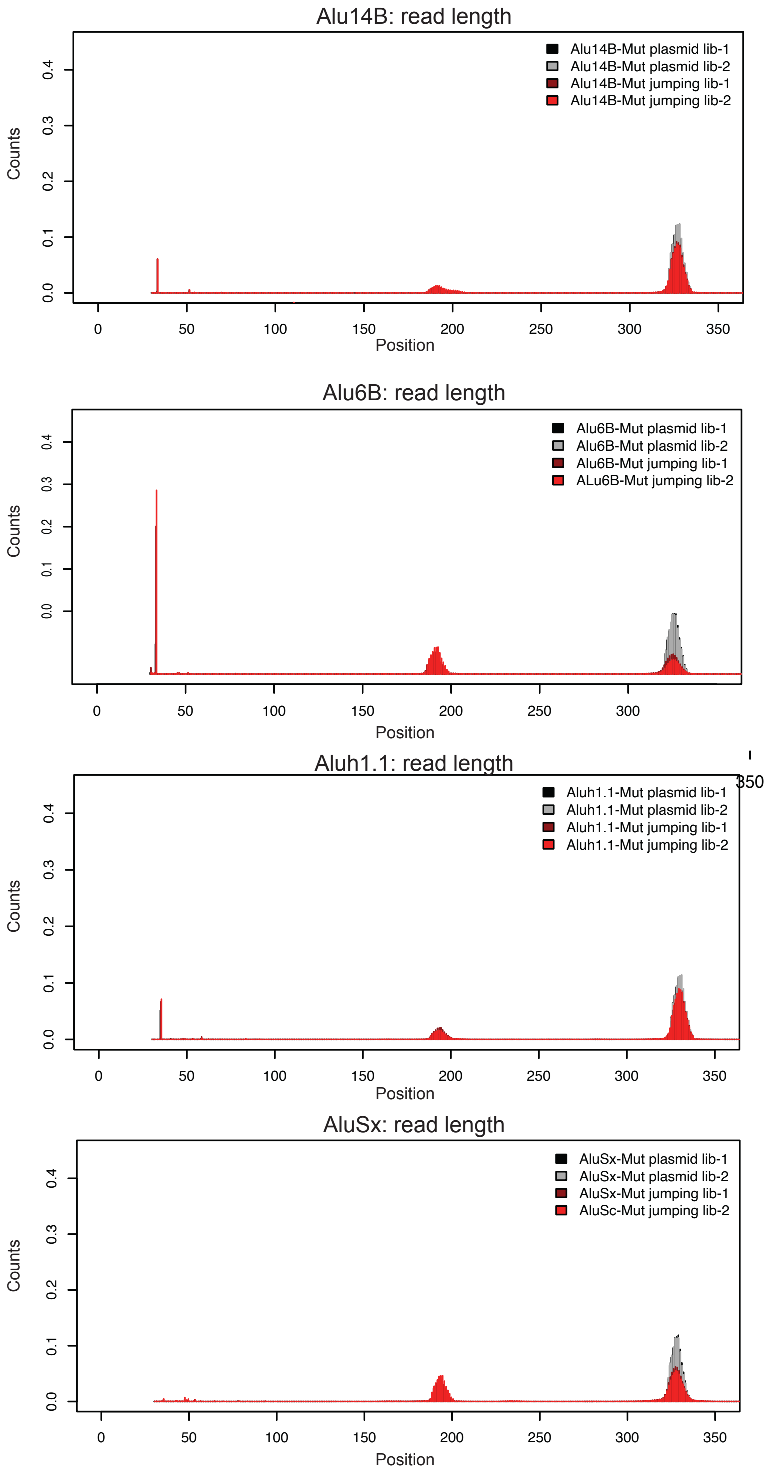


**Extended Data Fig. 7: Proportion of read count and read length in each library in both plasmid and jumping libraries.**
Read counts on Y axis and nucleotide position on Alu on X-axis calculated by MD-Mismatching positions/bases that records positional information from the reference sequence. Both plasmid and jumping library info in overlayed. A small subset of reads found in the jumping libraries were shorter (approx. 200bp) and missing the right monomer arm of the Alu element.

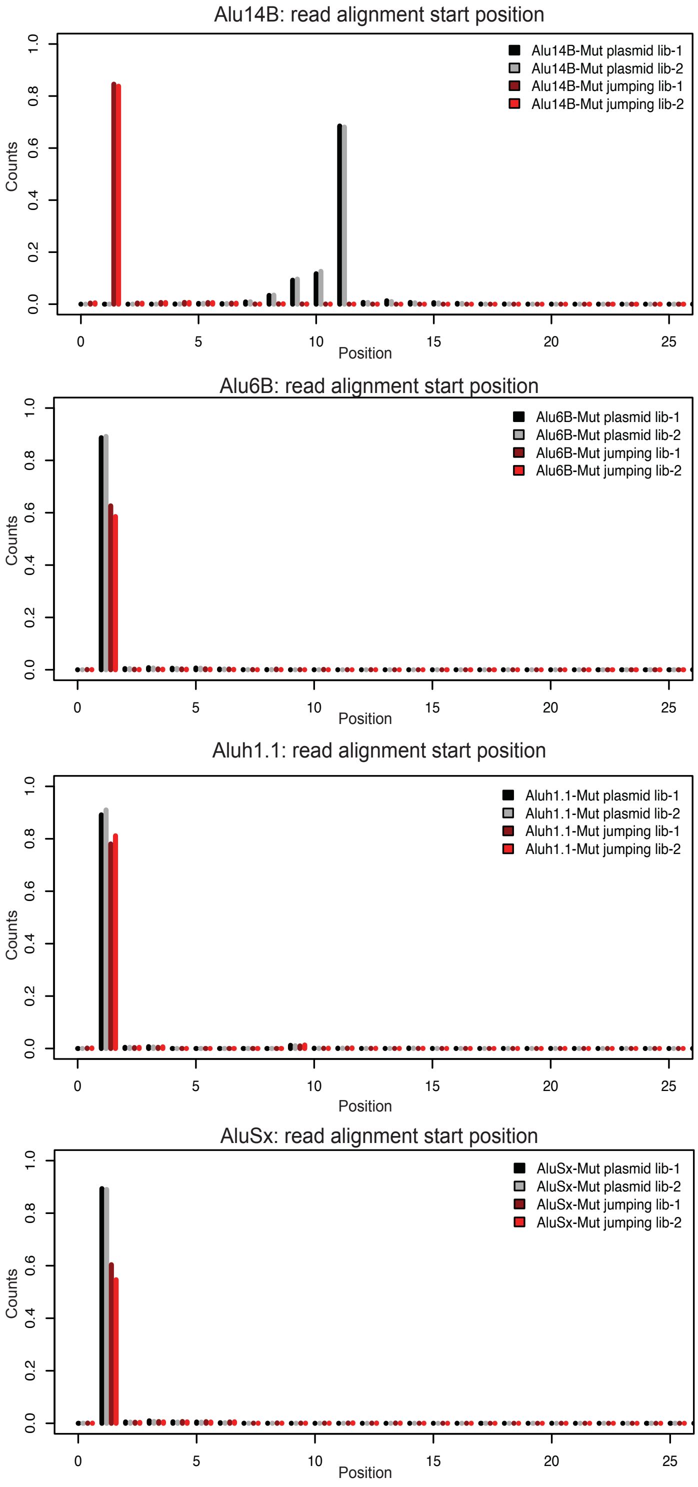


**Extended Data Fig. 8: Proportion of read count in the start position for each library in both plasmid and jumping libraries.**
Read counts on Y axis and nucleotide position on Alu on X-axis calculated by MD-Mismatching positions/bases that records positional information from the reference sequence for the 25 positions of the reference Alu sequence.

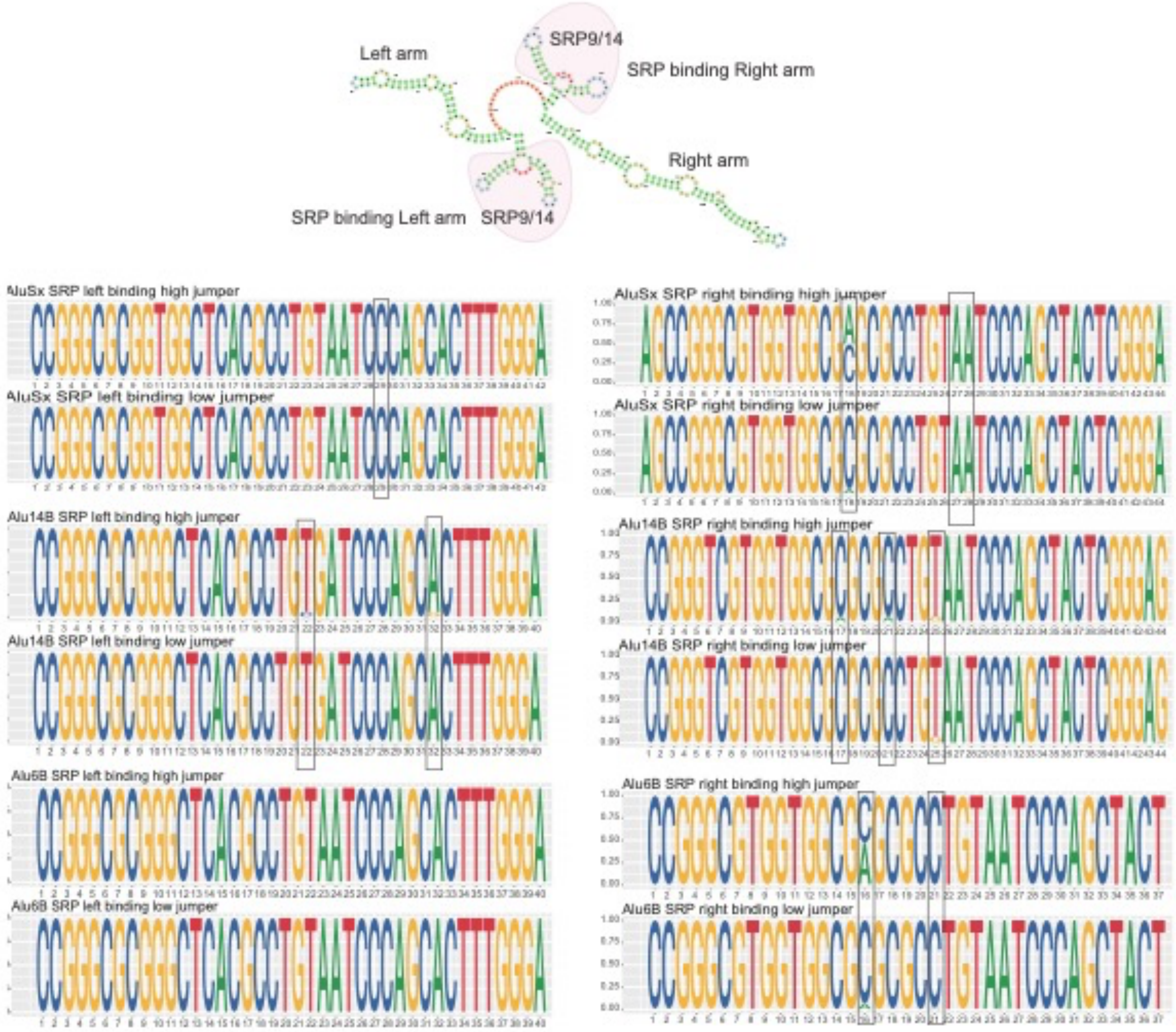


**Extended Data Fig. 9: Secondary structure of Alu RNA.**
Secondary structure of Alu RNA with the left and right SRP binding domain motifs in the left and right panel respectively. The top panel of each Alu shows the motif probability in high jumpers and the bottom panel shows the motif probability in low jumpers with some of the notable changes in nucleotide frequencies highlighted with a black rectangle.
